# A baseline for the genetic stock identification of Atlantic herring, *Clupea harengus*, in ICES Divisions 6.a, 7.b–c

**DOI:** 10.1101/2022.04.07.487341

**Authors:** Edward D. Farrell, Leif Andersson, Dorte Bekkevold, Neil Campbell, Jens Carlsson, Maurice W. Clarke, Afra Egan, Arild Folkvord, Michaël Gras, Susan Mærsk Lusseau, Steven Mackinson, Cormac Nolan, Steven O’Connell, Michael O’Malley, Martin Pastoors, Mats E. Pettersson, Emma White

## Abstract

Atlantic herring in ICES Divisions 6.a, 7.b-c comprises at least three populations, distinguished by temporal and spatial differences in spawning, which have until recently been managed as two stocks defined by geographic delineators. Outside of spawning the populations form mixed aggregations, which are the subject of acoustic surveys. The inability to distinguish the populations has prevented the development of separate survey indices and separate stock assessments. A panel of 45 SNPs, derived from whole genome sequencing, were used to genotype 3,480 baseline spawning samples (2014-2021). A temporally stable baseline comprising 2,316 herring from populations known to inhabit Division 6.a was used to develop a genetic assignment method, with a self-assignment accuracy >90%. The long-term temporal stability of the assignment model was validated by assigning archive (2003-2004) baseline samples (270 individuals) with a high level of accuracy. Assignment of non-baseline samples (1,514 individuals) from Division 6.a, 7.b-c indicated previously unrecognised levels of mixing of populations outside of the spawning season. The genetic markers and assignment models presented constitute a ‘toolbox’ that can be used for the assignment of herring caught in mixed survey and commercial catches in Division 6.a into their population of origin with a high level of accuracy.

## Introduction

Fish stock identification has been an important prerequisite for fisheries stock assessment throughout its history (Cadrin & Secor, 2009). However, the central fundamental weakness that remains in many existing stock assessments is the inaccurate recognition, definition and delineation of ‘stocks’ for data collection and aggregation. Traditionally, exploited stocks have been defined, assessed and managed according to geographical and political features or regions. Such is the case in the northeast Atlantic (FAO Major Fishing Area 27) where the European Union (EU) defines the term ’stock’ as ‘a marine biological resource that occurs in a given management area’ and delineates and names stocks using ICES (International Council for Exploration of the Sea) Statistical Areas (Anon, 2014). As more information becomes available, it is evident that the temporal and spatial distributions of most fisheries resources are not aligned to these artificial divisions (Kerr *et al*., 2016) and that biological populations are more dynamic and complex (Reiss *et al*., 2009; Stephenson, 2002).

Whilst delineation by predefined area may be convenient for management and regulation purposes, accurately assessing the status, biomass and sustainable exploitation rates of mixed ‘stocks’ is inherently difficult if not impossible, as they do not correspond to biological units. Fisheries dependent and independent data may be confounded, which may mask changes in the abundance of individual populations and lead to biased estimates of population abundance and overexploitation of smaller populations (Hintzen *et al*., 2015). It is thus critical to identify the underlying population structure of fisheries resources in order to identify the appropriate level at which to aggregate or segregate data for defining assessment and management units. It is also important to be able to assign individuals in mixed survey and commercial catches to the population or assessment unit to which they belong (Casey *et al*., 2016; Hintzen *et al*., 2015) in order to ensure the validity of data for inclusion in stock specific assessments. An ideal method of stock identification should be reproducible among laboratories and enable monitoring of the spatial and temporal integrity of a stock.

There is a long history of research into the characterisation of Atlantic herring (*Clupea harengus* Linnaeus, 1758) populations using a wide variety of different techniques, including life-history characteristics, morphometric and meristic characters of whole bodies and otoliths, parasite analyses, physical tagging and genetic approaches (see Farrell *et al*., 2021; Hatfield *et al*., 2005; McQuinn, 1997). Whilst many of the approaches have purported to offer reliable methods of discrimination between different populations, the reality is that confusion surrounding the population structure in herring across its distribution has persisted. This has prevented the identification of populations and hampered the delineation of stocks in many cases, for instance in the waters around Ireland and Britain where ICES currently assesses five herring stocks. The North Sea autumn spawning stock (ICES subarea 4, Divisions 3.a and 7.d.) is the most abundant and well-studied (Saville and Bailey, 1980) and is considered to be a complex of four spawning components (the autumn spawning Shetland/Orkney, Buchan, Banks components and the winter spawning Downs component), which are largely managed as one unit (Dickey-Collas *et al*., 2010; Simmonds, 2009). The definition of the western herring stocks has changed considerably over the last five decades (see Farrell *et al*., 2021; ICES, 2015) and the main stocks are currently recognised as: 6.a.N (*6aN_Aut*); 6.a.S, 7.b and 7.c (*6aS*); Division 7.a North of 52°30’N (Irish Sea/*IS*); Divisions 7.a South of 52°30’N, 7.g, 7.h, 7.j and 7.k (Irish Sea, Celtic Sea, and southwest of Ireland, *CS*) (Figure 1; ICES, 2014). The *6aN_Aut* herring spawn in Autumn (Sept/Oct) off Cape Wrath on the north coast of Scotland, the *6aS* herring spawn in winter (Nov-Feb) primarily off the coast of Donegal in the northwest of Ireland, *IS* herring spawn in Autumn (Sept/Oct) mainly on the Douglas Bank east of the Isle of Man in the Irish Sea and Celtic Sea herring spawn in winter (Nov-Feb) off the south coast of Ireland. Several groups of spring spawning (Feb-May) herring are also known to occur in the Minch (*6aN_Sp*), Clyde and Milford Haven, though these are not currently assessed and are believed to be small populations (see review in Farrell *et al*., 2021). Other autumn/winter spawning herring groups are also found in the western English Channel and Bristol Channel (ICES Divisions 7.e and 7.f, respectively), though no assessment is made of these groups and there are no management measures in place.

**Figure 1.**
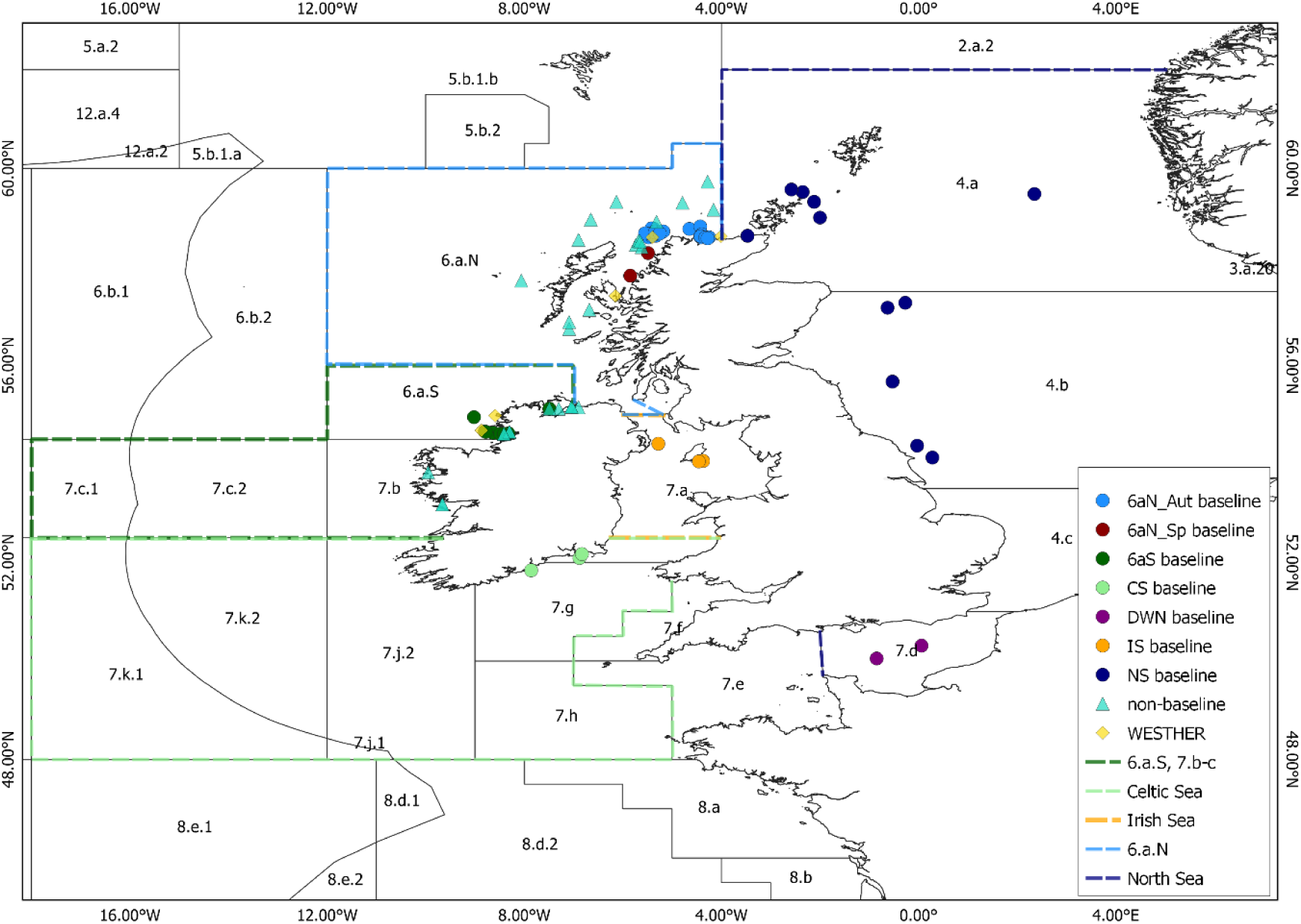
The distribution of herring samples collected and analysed in the current study. The sample type and current stock boundaries are indicated according to the legend.

The stock divisions for herring assessments and management around Ireland and Britain are largely based on the recognition of temporal and spatial differences in spawning season and grounds and are believed to broadly align with biological population structure (ICES, 2015). Though some geographic and political boundaries are still in place, the mixing across these boundaries is unclear. This is evident in ICES Division 6.a, where the *6aN_Aut* stock is separated from the North Sea autumn spawning stock by the 4° west line of longitude, despite there being no biological evidence that these represent different populations (see review in Farrell *et al*., 2021). Within Division 6.a the herring are subdivided into two stocks (Figure 1) by the 56° north line of latitude and 7° west line of longitude (ICES, 1982). Herring caught or surveyed to the north or east of this boundary (excluding the Clyde area) are included as part of the *6aN_Aut* stock regardless of their population of origin or their spawning time. This includes herring caught in Lough Foyle in NW Ireland, whose waters are bisected by the 7° west line. Most of Lough Foyle is west of this line, however the mouth of Lough Foyle is east of this line and hence the herring in the whole Lough are considered to be part of the *6aN_Aut* stock despite having no affinity to this population. Herring caught to the south and west of the 56° and 7° lines are considered to be part of the *6aS* stock in combination with herring in Divisions 7.b and 7.c. Adult herring from different populations, both within Division 6.a (*6aN_Aut*, *6aS*, *6aN_Sp*) and possibly from adjacent populations (*IS* and *CS*) are believed to form mixed aggregations on common feeding grounds in Division 6.a during summer (Hatfield *et al*., 2005). It is during this time that they are surveyed by the annual Malin Shelf Herring Acoustic Survey (MSHAS), part of the internationally coordinated Herring Acoustic Survey (HERAS), which is the primary tuning index used in the stock assessments of Division 6.a herring. The inability to assign herring catches from the MSHAS into their population of origin prevents the development of separate indices of abundance for the populations in Division 6.a, therefore ICES has conducted a combined assessment of these populations since 2015 (ICES, 2015), which provides combined management advice. Combined management of separate stocks can only be precautionary if the two stocks are of similar size and are homogeneously distributed together in commercial catches. If these conditions are not met, uncertainty of the status of each of the individual stocks increases, as does the risk that one stock may sustain higher fishing mortality than the other.

Genetic assignment methods, which compare genetic data from individuals to genetic profiles of reference samples from potential source populations to determine population of origin (Manel *et al*., 2005), offer the potential to resolve these issues. However, the incorporation of genetic assignment methods into regular fisheries data collection, assessment and management has been slow (Bernatchez *et al*., 2017; Reiss *et al*., 2009; Waples *et al*., 2008), as many existing genetic studies have been hampered by high cost, inadequate sampling coverage, low numbers of suitable molecular markers and low power to detect genetic structure. The advent of high-throughput sequencing (HTS) technologies fundamentally changed the way in which genetic sequence data are generated (see Hemmer-Hansen *et al*., 2014; Davey *et al*., 2010). It is now possible to generate large genomic data sets for non-model species, which facilitate the identification of genetic loci with high discriminatory power for resolving specific population differentiation questions (Martínez Barrio *et al*., 2016; Nielsen *et al*., 2012). There has also been a shift toward the analysis of sequence variation of functional, adaptive significance rather than just neutral DNA sequence variation (Mariani & Bekkevold, 2013). This approach focuses on identifying adaptive markers that are under diversifying selection and may reflect distinctive features of local populations (Nielsen *et al*., 2012). Small panels of high-graded markers may be selected to develop efficient and cost-effective genetic assignment tools for informing marine fisheries assessment and management (Hemmer-Hansen *et al*., 2018).

Assignment methods that attempt to solve classification problems rely on computing a discriminant function based on samples from potential source populations and then classify unknown individuals to the group with the highest discriminant score (Manel *et al*., 2005). Genetic assignment methods have traditionally relied on using the genotypic frequency distribution under the assumption of Hardy–Weinberg equilibrium (HWE) and linkage equilibrium in each source population as their discriminant function (Manel *et al*., 2005). These genetic assignment methods can be broadly divided into Bayesian (Rannala and Mountain, 1997), frequency (Paetkau *et al*., 1995) and distance based (Cornuet *et al*, 1999) methods (Hauser *et al*., 2006). The underlying assumptions of the methods are quite similar although the distance-based methods may be less sensitive to violations of population genetic expectations such as HWE and linkage equilibrium (Cornuet *et al*., 1999). These methods are commonly implemented in the software GeneClass2 (Piry *et al*., 2004). In the absence of baseline data to guide classification, Bayesian clustering methods may be used to delineate clusters of individuals based on their multi-locus genotypes and assign individuals to their individual clusters (Manel *et al*., 2005). However, these Bayesian clustering analyses such as that implemented in the software *Structure* (Pritchard *et al*., 2000) are also constrained by the underlying assumptions of HWE and linkage equilibrium. Multivariate analysis has several advantages over other classical approaches used in population genetics and genetic assignment, the foremost of which is that they do not require the assumptions of HWE or linkage equilibrium (Jombart *et al*., 2009). Multivariate approaches are particularly suited to solving classification problems when used in the form of supervised machine learning (SML) approaches. SML is concerned with predicting the value of a response label/category on the basis of the input variables/features (Schrider and Kern, 2018). When empirical data are available, SML trains an algorithm based on a training set of the labelled data, which can then be used to predict the category of unknown data. Support Vector Machines (SVM) are a set of SML methods that can be used for classification problems. The objective of SVM algorithms is to find a hyperplane in an N-dimensional space (N - the number of features) that distinctly classifies the data point (see James *et al*., 2013). SVM models can also be used to classify non-linear data through use of non-linear kernels (James *et al*., 2013) and can be optimised by adjusting parameters, including cost and gamma, which control the stringency of the boundary and the influence of single training datapoints, respectively. The *R* package *assignPOP* (Chen *et al*., 2018) has recently made the use of SVM models for assignment more accessible and also allows for the integration of genetic and non-genetic data within the same model, which is an advantage in many stock identification studies which also collect morphometric data.

Recent studies utilising Whole Genome Sequencing (WGS) approaches, have comprehensively clarified the genetic population structure of Atlantic herring across its distribution and have illustrated that herring populations show strong population structure inferred to be associated with a high level of local ecological adaptation (Han *et al*, 2020; Lamichhaney *et al*., 2017; Martinez Barrio *et al*., 2016). Genetic markers associated with loci under selection have also been proven to provide a significantly better resolution to distinguish population structure than neutral genetic markers (Han *et al*., 2020). From the circa 10 million Single Nucleotide Polymorphisms (SNPs) identified by Han *et al*. (2020) a small subset of circa 800 SNPs, associated with ecological adaptation to different geographic areas and spawning conditions, were shown to be able to discriminate all the sampled populations of herring from across the species distribution. The populations around Ireland and Britain are the southernmost ecomorphs of herring in the Northeast Atlantic and are genetically distinct from the other adjacent Northeast Atlantic herring populations, including Norwegian Spring Spawning herring to the north and the Baltic herring to the east, from which they can be discriminated with a small subset of 12 independent loci (Han *et al*., 2020). The populations sampled around Ireland and Britain could be further subdivided into four main groups: the spring spawning herring from the Minch (*6aN_Sp*) and the Clyde; the *6aN_Aut* and North Sea autumn spawning herring, which were indistinguishable from each other, thus supporting the conclusion these stocks are most likely a single population; the *6aS* herring, which were differentiated from all populations but more closely related to the southern group consisting of *CS*, *IS* and Downs herring, which had the lowest level of genetic differentiation between them. Identification of the primary genome level differences between the herring populations around Ireland and Britain offers the potential to develop a genetic assignment method for discriminating and resolving the outstanding issues of separating mixed survey and commercial catches in ICES Divisions 6.a, 7.b**–**c.

The aims of the current study were therefore to:

(i) validate a small ‘toolbox’ of genetic markers, from those identified by Han *et al*. (2020), that could be used to develop genetic baselines for the individual populations in Divisions 6.a, 7.b**–**c.
(ii) develop a temporally stable genetic baseline dataset by collecting and analysing multiple years of spawning baseline samples from each population.
(iii) develop, test and validate an assignment model for the genetic assignment of individuals of unknown origin collected in Divisions 6.a, 7.b**–**c back to their population of origin.

## Materials and Methods

### Sampling and DNA isolation

Samples of herring were collected from the catches of fisheries surveys and commercial fishing operations, between 2014 and 2021 in the core ICES Divisions 6.a, 7.b**–**c area and on the adjacent populations where possible. Each fish was measured for total length (to the 0.5 cm below), total body weight to the nearest 1 g and assessed for sex and maturity. Samples processed by Marine Scotland Science (MSS) were maturity staged using the 9-point scale, those processed by the Irish Marine Institute (MI) were maturity staged using the 8-point scale and samples processed by the Wageningen University and Research (WUR) on behalf of the Dutch Pelagic Freezer Trawler Association (PFA) were maturity staged using the ICES 6-point scale (ICES, 2011). All maturity stages were converted to the ICES 6-point scale according to Mackinson *et al*. (2021). A 0.5 cm^3^ piece of tissue was excised from the dorsal musculature of each specimen, taking care to avoid skin and scales, and stored in absolute ethanol at 4°C. Archive fin clips were also available from the spawning baseline samples collected during the WESTHER project 2003-2004 (FP5-LIFE QUALITY Q5RS-2002-01056; Hatfield *et al*., 2005). Total genomic DNA (gDNA) was extracted from c.10 mg of tissue or fin clip from each fish using 300 µl of 10% Chelex suspension and 5 µl of Proteinase K (20 mg/µl). Extracted DNA was stored in 96 well PCR plates at -20°C until ready for genotyping.

### Genetic marker identification

The SNPs used in the current study (ESM Table 1) were identified during the GENSINC project (GENetic adaptations underlying population Structure IN herring; Research Council of Norway project 254774) and were derived from the analyses of WGS of pooled samples from herring populations across the species distribution, which was undertaken to study the biological significance of the genetic variants underlying ecological adaptation in the Atlantic herring (Han *et al*., 2020; Lamichhaney *et al*., 2012; Martinez Barrio *et al*., 2016; Pettersson *et al*., 2019). The subset of SNPs was selected following testing of candidate SNPs with the highest delta allele frequency (dAF) values from the major genomic regions of divergence in the contrasts between populations around Ireland and Britain. The 45 SNPs selected were distributed across 8 chromosomes and comprised 14 loci (ESM Table 1). Linked SNPs were retained in the panel in order to add a level of redundancy and ensure that key genomic regions (loci) were well represented even in the instance of missing genotype data from an individual.

**Table 1.** Pairwise multi-locus *F_ST_* (above the diagonal) for the *baseline dataset* and associated *P*- values (below the diagonal) with the temporal replicates condensed.

|  | 6aS | CS | IS | 6aN_Aut | 6aN_Sp | NS | DWN |
| --- | --- | --- | --- | --- | --- | --- | --- |
| 6aS |  | 0.09 | 0.13 | 0.20 | 0.35 | 0.24 | 0.12 |
| CS | 0.0001 |  | 0.08 | 0.23 | 0.64 | 0.32 | 0.01 |
| IS | 0.0001 | 0.0001 |  | 0.20 | 0.67 | 0.29 | 0.08 |
| 6aN_Aut | 0.0001 | 0.0001 | 0.0001 |  | 0.57 | 0.02 | 0.24 |
| 6aN_Sp | 0.0001 | 0.0001 | 0.0001 | 0.0001 |  | 0.60 | 0.68 |
| NS | 0.0001 | 0.0001 | 0.0001 | 0.0001 | 0.0001 |  | 0.33 |
| DWN | 0.0001 | 0.0001 | 0.0001 | 0.0001 | 0.0001 | 0.0001 |  |

### Genotyping

The majority of samples were genotyped utilising a genotyping by sequencing approach (Vartia *et al*., 2016) described in detail and validated in Farrell *et al*. (2016; 2021). In short, locus-specific forward and reverse primers were designed for SNP loci with the Primer3 application (Rozen and Skaletsky, 2000) in Geneious® 7.0 (Kearse *et al*., 2012) with optimal primer length set at 20bp and product size range at 120-180bp. Primers were designed to bind in conserved flanking regions to minimise the possibility of null alleles and were cross-referenced with existing genome sequence data to identify primers that annealed to multiple regions, which if detected were excluded. The forward and reverse locus-specific primers were adapted, to facilitate combinatorial barcoding of amplicons, by adding either an M13-R (5’-GGAAACAGCTATGACCAT-3’) or CAG (5’-CAGTCGGGCGTCATCA-3’) universal tail to the 5’ end and were divided into multiplex panels in MultiPLX 2.1 (Kaplinski *et al*., 2005). A set of ninety-six 11bp combinatorial barcodes were used to identify individuals within pooled sequencing runs. An M13-R universal tail was added to the 3’ end of forty-eight of the barcodes and a CAG universal tail to the 3’ end of the remaining forty-eight barcodes, yielding 2,304 possible combinations. The tagged primers and tagged barcodes were tested for the formation of secondary structures (hairpins, primer dimers and hetero dimers) with the IDT OligoAnalyzer Tool 3.1 (http://eu.idtdna.com/calc/analyzer).

Amplification and barcoding reactions were carried out using a two-step PCR as described in Farrell *et al*. (2016 and 2021). In short, the first PCR involved the amplification of the target SNPs and the second PCR involved the incorporation of the combinatorial barcodes for individual identification. Following PCR amplification each plate of amplicons was pooled and then standardised for concentration and combined into a single sample before sending for library preparation and amplicon sequencing by a third-party sequencing service provider. Six different herring amplicon sequencing runs were conducted over the course of the current project using both the Illumina MiSeq and HiSeq platforms. The raw data from these runs were treated following the same protocols in order to derive the final individual genotypes. Raw FASTQ sequence data were downloaded from Illumina BaseSpace and initial quality control was performed using FastQC (Babraham, 2016). Reads were sorted and grouped using a modified python script (Vartia *et al*., 2016) based on the Levenshtein distance metric. The raw sequence data were processed by identifying sequence reads containing the forward and reverse combinatorial barcodes and the locus-specific primers. Reads were sorted hierarchically and grouped into five separate FASTA files as reads with: no barcode, one barcode, two barcodes and no primers, two barcodes and two non-matching primers, two barcodes and two matching primers. Only reads containing two barcodes and two matching primers were included in further analyses. These reads were grouped by locus and individual before removing the barcode from the sequences.

SNP genotyping was automated by using a modified Perl script from the Genotyping-in-Thousands by sequencing (GT-seq) approach (Campbell *et al*., 2015), which counts amplicon-specific sequences for each allele, and uses allele ratios to determine the genotypes. The Perl scripts were modified in the current project to use the output of the custom python scripts as the input. The default settings of the GT-Seq Perl script designated allele ratios >10.0 to be called as homozygous for allele 1, ratios <0.1 to be called as homozygous for allele 2, and ratios between 0.2 and 5.0 to be called as heterozygous (Campbell *et al*., 2015). These ratios were optimised for the data and markers in the current study by analysing each marker separately and plotting the genotyping calls from which new ratios were calculated for each marker. The average designated allele ratios in the current study were >5.0 to be called as homozygous for allele 1, ratios <0.2 to be called as homozygous for allele 2, ratios between 0.3 and 3.33 to be called as heterozygous and ratios between 3.34-4.9 and 0.201-0.29 were called as NA (NA = no genotype call made). Individuals with less than 10 reads at a particular locus were also designated as NA. Only individuals with greater than 89% genotyping success (i.e. 40/45 genotypes) were retained in the dataset.

Genotyping of the majority of samples collected from quarter three 2019 to 2021 was undertaken by a commercial provider; IdentiGEN, Dublin, Ireland, using their proprietary *IdentiSNP* genotyping assay chemistry, which utilises target specific primers and universal hydrolysis probes. Following an end- point PCR reaction, different genotypes were detected using a fluorescence reader. Concordance between the two genotyping methods was confirmed by genotyping a subset of samples from each of the target populations (n=24 per population) and confirming that the same genotypes were called with each method (data not shown).

### Baseline dataset analyses

It should be noted that the aim of the current study was not to undertake an exhaustive population genetics and demographic study of the herring populations around Ireland and Britain but was to develop a genetic based method to separate the herring caught in putatively mixed survey and commercial catches in ICES Divisions 6.a, 7.b and 7.c into their population of origin. The analytical approaches followed were tailored to this specific task. The limited number of genetic markers used in the current study were high graded to maximise the power of discrimination between the core Division 6.a populations and in some instances comprised multiple SNPs from a small number of loci. Therefore, the dataset may not be suitable for conventional population genetic analyses and as such some of the analyses presented (e.g. estimation of fixation indices) were for exploratory purposes only.

Deviations from Hardy–Weinberg equilibrium (HWE), linkage disequilibrium (LD), and excess and deficiency of heterozygotes in the *full baseline dataset* (see Results) were tested with Genepop 4.2 using default settings (Rousset, 2008). Microsatellite Analyzer (MSA) 4.05 (Dieringer and Schlötterer, 2003) was used, under default settings, to assess multi-locus pairwise *F_ST_* with 1,000 bootstrap replications and 10,000 permutations. In all cases with multiple tests, significance levels were adjusted using the sequential Bonferroni technique (Rice, 1989). In order to visualise the pairwise *F_ST_* results and to explore the relationships between the different samples, Principal Coordinate Analysis (PCoA) using the covariance standardised method was conducted in GenAlEx 6.51b2 (Peakall and Smouse, 2012).

Discriminant analysis of Principal Components (DAPC), from the *R* package *adegenet* (Jombart, 2008; Jombart *et al*., 2010), is a multivariate approach that transforms multi-locus genotype data using PCA to derive a set of uncorrelated variables, which serve as input for discriminant analysis (DA). The DA aims to maximize among-group variation and minimize within-group variation. DAPC does not make assumptions of underlying population genetic processes (e.g. neutrality, linkage equilibrium, Hardy– Weinberg equilibrium), therefore it was appropriate to use this approach with the data in the current study. In the first instance DAPC was run using the 64 baseline samples as the input groups and retaining all PCs and discriminant functions. The DAPC was run again after the temporal samples were combined to form seven groups (*6aS*, *CS*, *IS*, *6aN_Aut*, *6aN_Sp*, *NS*, *DWN*) which represented the putative populations in the study area. Following further analyses (see Results) a reduced *6a baseline dataset* consisting only of samples from groups that are confirmed as being present in Division 6.a i.e. *6aS*, *6aN_Aut* and *6aN_Sp,* was analysed using DAPC. In this instance DAPC was conducted as before with prior definition of group membership and also following a second approach using the *find.clusters* function to infer genetic clusters. This function transforms the data using principal component analysis (PCA), then runs the *K*-means algorithm (function *kmeans* from the *stats* package) with increasing values of *K* and computes Bayesian Information Criterion (BIC) to assess the best supported model.

### Assignment model development

The *R* package *assignPOP* (Chen *et al*., 2018), which performs population assignment using a machine-learning framework, was used to develop the assignment model. *assignPOP* uses Monte-Carlo cross-validation (*assign.MC*) to divide the baseline data into a training dataset and test dataset. The assignment model is developed with the training dataset and subsequently tested with the independent test dataset, which avoids introducing ‘high-grading bias’ (see Anderson, 2010). As the Monte-Carlo procedure samples random individuals each time, it does not guarantee that every individual is sampled. Therefore, *assignPOP* can perform an additional method of *K*-fold cross-validation (*assign.kfold*), which involves randomly dividing the individuals from each population into *K* groups and then using one group from each population as test individuals and the remaining *K-1* groups as the training individuals. Assignment tests are performed until every group and hence individual is tested, resulting in *K* tests. *assignPOP* has a number of classification model options including the *SVM* model from the *R* package *e1071* (Meyer *et al*., 2015). Based on the results of the aforementioned baseline dataset analyses it was decided to develop the assignment model in *assignPOP* trained on the reduced *6a baseline dataset* as the constituent populations had the highest level of discrimination between them with the current marker panel. As per the DAPC analyses, the assignment model was developed using two different approaches with, in this case, each approach conducted at two hierarchical levels.

*Approach 1* used the *6a baseline dataset* with the predefined *6aS*, *6aN_Aut* and *6aN_Sp* population groups. The assignment was conducted at two hierarchical levels based on the power to discriminate the different groups in the DAPC analyses. In *Level 1 6aS* and *6aN_Sp* were combined and tested against *6aN_Aut*. In *Level 2* the combined *6aS/6aN_Sp* group was split, and the individual groups tested against each other. *Approach 2* was also performed in a hierarchical manner as per *Approach 1*. However, *Approach 2* was initially independent of the assumptions of prior populations and instead used the output of the *K*-means clustering analyses of the *6a baseline dataset* to identify different baseline assignment clusters. In the cases where multiple clusters represented a single assumed population of origin these clusters were combined.

In order to avoid over-fitting, the model and to objectively determine the optimum number of PCs to be used in both assignment approaches, DAPC cross-validation was conducted with the *xvalDapc* function in *adegenet*. Exploratory analyses were conducted in *assignPOP* to determine the optimum model and kernel for the assignment model and the *tune*, *tune.control* and *best.svm* functions in *R* package *e1071* (Meyer *et al*., 2015) were used to perform a grid search for the optimum values for cost and gamma. These parameters were used for testing the rate of self-assignment using both Monte-Carlo and *K*-fold cross-validation to estimate membership probability. In order to avoid unbalanced sample sizes among the baseline groups the number of individuals in the training sets were specified and were limited by the number of individuals in the smallest group. *Level 1* assignments were tested with 200, 400, 600 and 800 individuals in the training set, whilst the *Approach 1-Level 2* assignment was tested with 50 and 75 individuals and *Approach 2-Level 2* assignment with 100, 150 and 200 individuals. Both Monte-Carlo and *K*-fold cross-validation were performed using 25%, 50%, 75% and 100% of the highest *F_ST_* loci (*loci.sample=“fst”*) and all tests were conducted with 100 iterations.

An important consideration when developing the assignment model was to determine how many genetic markers were required for accurate assignment using either of the approaches and at either of the levels. This enabled the threshold for missing data of unknown samples to be set with a robust basis without compromising the integrity of the assignments. In order to do this the Monte-Carlo cross validation analyses were run again with random sampling of loci (*loci.sample=”random”*) rather than highest *F_ST_* loci and were run with 20-100% of loci in 10% intervals. All other parameters were the same as the previous runs.

### Assignment model validation with archive samples

As an additional validation of the baseline assignment models the WESTHER baseline samples collected in 2003/2004 in Division 6.a were used as known-unknown samples and assigned to the contemporary baseline in order to test the long-term temporal stability of the assignment models. The WESTHER samples were processed and genotyped following the same method as the other samples in the study. The assignments were conducted using the *assign.X* function in *assignPOP* (Chen *et al*., 2018) using the two hierarchical approaches and with the same model parameters as described above. For each approach, the *Level 1* assignment was conducted with all the individuals in the sample and the *Level 2* assignment, on a subset of individuals that required further assignment. A successful assignment probability threshold was set at 0.67, which indicated a situation where one assignment outcome was twice as likely as the alternate outcome. This was deemed an acceptable level of confidence given the high level of self-assignment accuracy of the baseline datasets. The final assignment call of each individual was based on a combination of the *Level 1* and *Level 2* assignments. Final sample assignments were plotted using the *draw.pie* function of the *R* package *mapplots*. In order to test the potential effect of increasing the assignment threshold and also to compare the relative assignment rates between the assignment approaches, the proportion of individuals falling below thresholds of 0.67, 0.7, 0.8 and 0.9 were also calculated.

### Exploratory analyses with contemporary non-baseline samples

Additional samples, which were not considered to be baseline samples (i.e. they were not collected on known spawning grounds or were not in the correct maturity stage) were also collected during the study. These non-baseline samples were used to further test the assignment model and also to provide an exploratory analysis of potential mixing of populations within Divisions 6.a, 7.b-c. The samples were assigned as per the WESTHER samples and divided by quarter for plotting using *mapplots*. As with the archive samples the effect of the range of assignment thresholds was also tested with these samples.

## Results

### Sampling and genotyping success rate

Due to the opportunistic nature of the sampling, the samples contained a significant mix of length classes and maturity stages (ESM Tables 2 and 3). In total 92 contemporary samples were collected (Figure 1 and ESM Table 2), comprising 6,591 individual herring of which 5,638 individuals (86%) passed the genotyping threshold of 89% (40/45 SNPs genotyped). All 45 SNPs were successfully genotyped in more than 92% of retained individuals. For the purposes of developing robust baselines for genetic assignment, it is critical to avoid including individuals with uncertain origin. Therefore, only samples with a significant number of maturity stage three (spawning) individuals (ICES, 2011), caught in close proximity to known spawning grounds at recognised spawning times were selected to be baseline samples. In order to further limit the potential for misclassification as a baseline spawning sample, only individuals classified as maturity stage three were included in the baseline dataset. The resulting contemporary *baseline dataset* contained 64 samples, comprising a total of 3,480 herring (Figure 1).

**Table 2.** Clustering analyses, using the *find.clusters* function in *adegenet*, of the *6a baseline dataset*. The percentage of each population group split by cluster and the percentage of each cluster split by population are shown.

| Populations by Cluster |  |  |  |
| --- | --- | --- | --- |
|  | 1+3+5 | 2 | 4+6 |
| 6aS | 3.5 | 13.8 | 82.7 |
| 6aN_Aut | 93.6 | 0.4 | 6.0 |
| 6aN_Sp | 2.0 | 95.1 | 2.9 |

| Clusters by Population |  |  |  |
| --- | --- | --- | --- |
|  | 1+3+5 | 2 | 4+6 |
| 6aS | 2.3 | 53.6 | 89.4 |
| 6aN_Aut | 97.5 | 2.3 | 10.5 |
| 6aN_Sp | 0.2 | 44.1 | 0.4 |

**Table 3.**
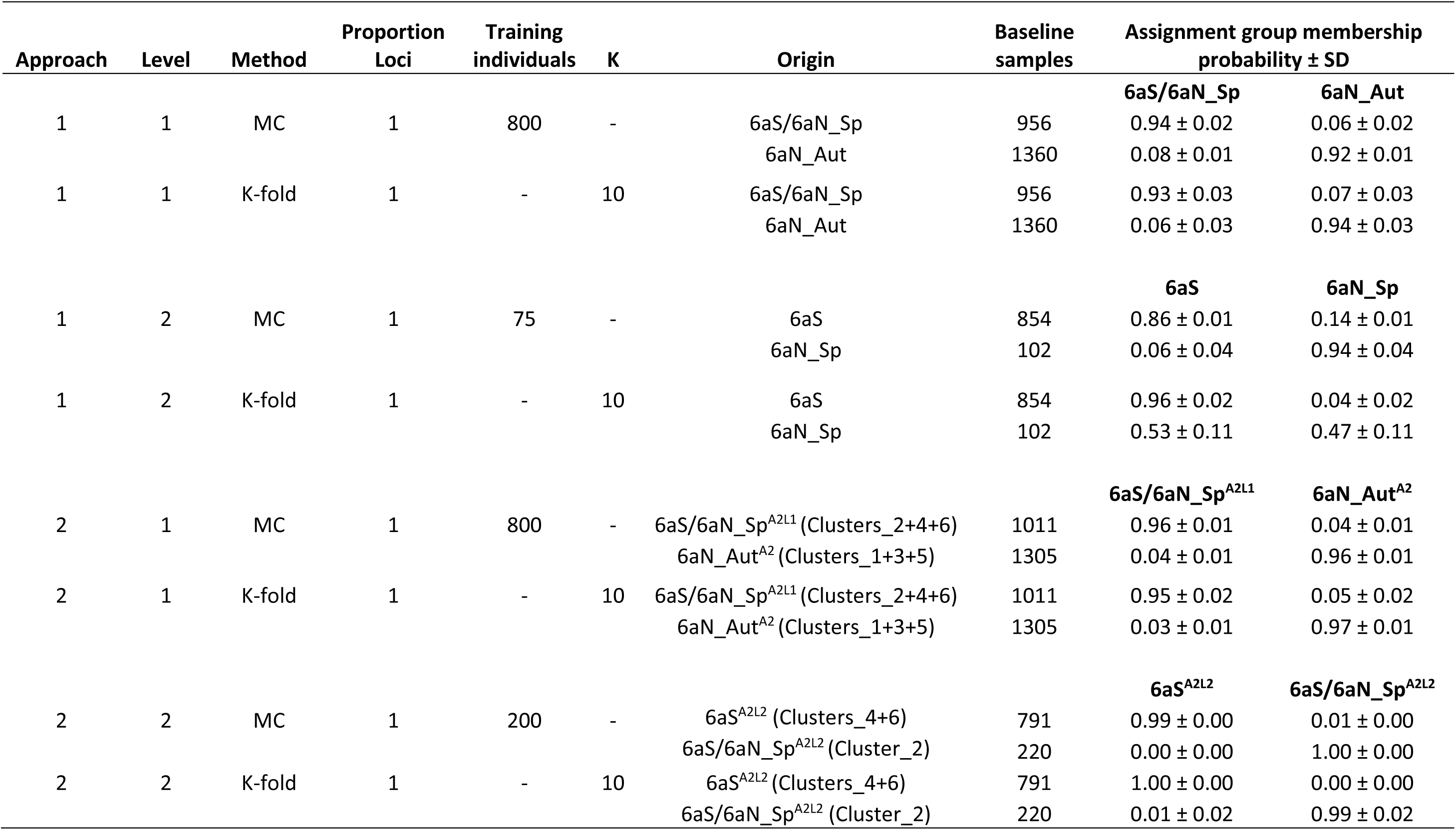
Monte-Carlo and *K*-fold cross validation results from the assignPOP analyses of the two assignment approaches.

The 28 remaining samples (1,514 individuals) were retained in a separate *non-baseline dataset*, to be used to test the assignment model and provide an exploratory analysis of potential mixing of populations within Divisions 6.a., 7.b-c. In addition, five archive baseline samples comprising 340 individuals collected in 2003 and 2004, were also available from the WESTHER project, of which 270 surpassed the genotyping quality control threshold. These samples were retained in an *archive dataset* for the purposes of having independent baseline samples for the validation of the assignment model and for testing the long-term temporal stability of the assignment model.

### Baseline dataset analyses

There were no significant patterns of deviation from HWE, heterozygote deficiency or heterozygote excess at the locus level (45 SNPs). At the population level significant deviations from HWE were observed in samples *6aN_Sp_18b* (10/45 SNPs), *6aS_17d* (11/45), *6aS_17e* (12/45), *6aS_19c* (6/45). Samples *6aS_17d* and *6aS_17e* also displayed indications of a significant heterozygote deficiency in eight and thirteen loci, respectively, which was likely the result of some of the *6aS* samples containing a mixture of early and later spawning components (see pairwise *F_ST_* results). Samples *6aN_Sp_18b*, *6aS_19c* and *DWN_18* displayed indications of significant heterozygote excess at ten, seven and seven loci, respectively. The significant indications of LD were in keeping with the loci already identified (ESM Table 1) from Han *et al*. (2020). All markers and all samples were retained in the *baseline dataset* for further analyses.

The analyses of multi-locus pairwise *F_ST_* (ESM Table 4) indicated proportionately higher *F_ST_*’s and significant differentiation between samples collected from the different putative populations except between the *6aN_Aut* and *NS* populations, which displayed little if any significant genetic differentiation among or between the temporal samples. There was no significant differentiation between the temporal samples from the *IS* or *DWN* population and there was little if any significant genetic differentiation between the temporal samples from the *CS* and *6aN_Sp*. There were some indications of differentiation between the temporal samples from *6aS*, with some of the samples collected in January and February (*6aS_17a*, *6aS_17b*, *6aS_18a*) showing a low level of differentiation from the other samples. The PCoA of the pairwise *F_ST_* results enabled a clearer interpretation and illustrated the clustering of samples within and between putative populations (Figure 2). The temporal samples from each population clustered together though some intrapopulation diversity was evident particularly among the *6aS*and *6aN_Aut* samples. The *6aN_Sp* samples were distinct from the *6aN_Aut* samples and were closely aligned with the late *6aS*samples i.e. those collected in quarter 1.

**Figure 2.**
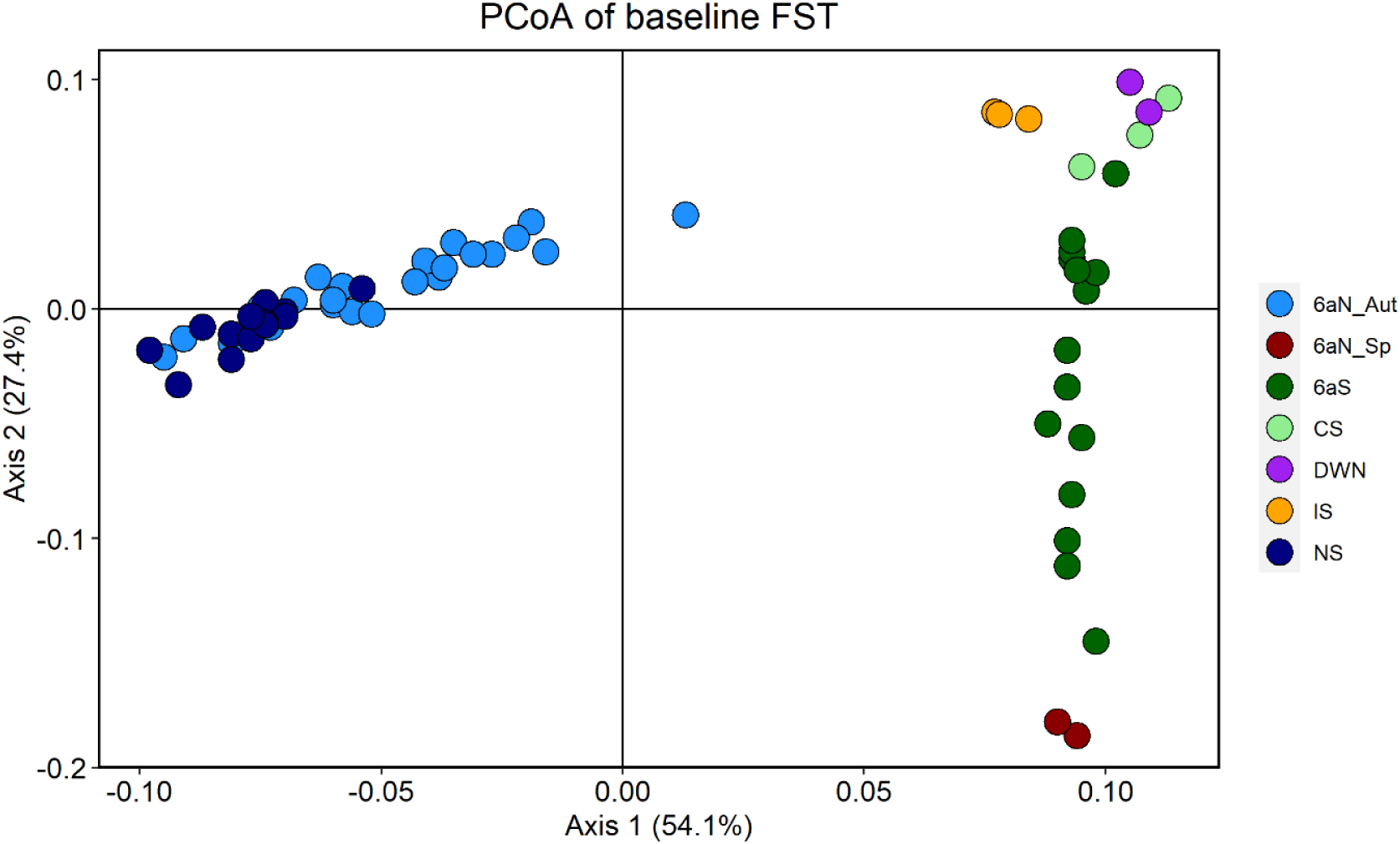
Principal Coordinate Analyses (PCoA) of the multi-locus pairwise *F_ST_* analyses of the baseline samples.

**Table 4.**
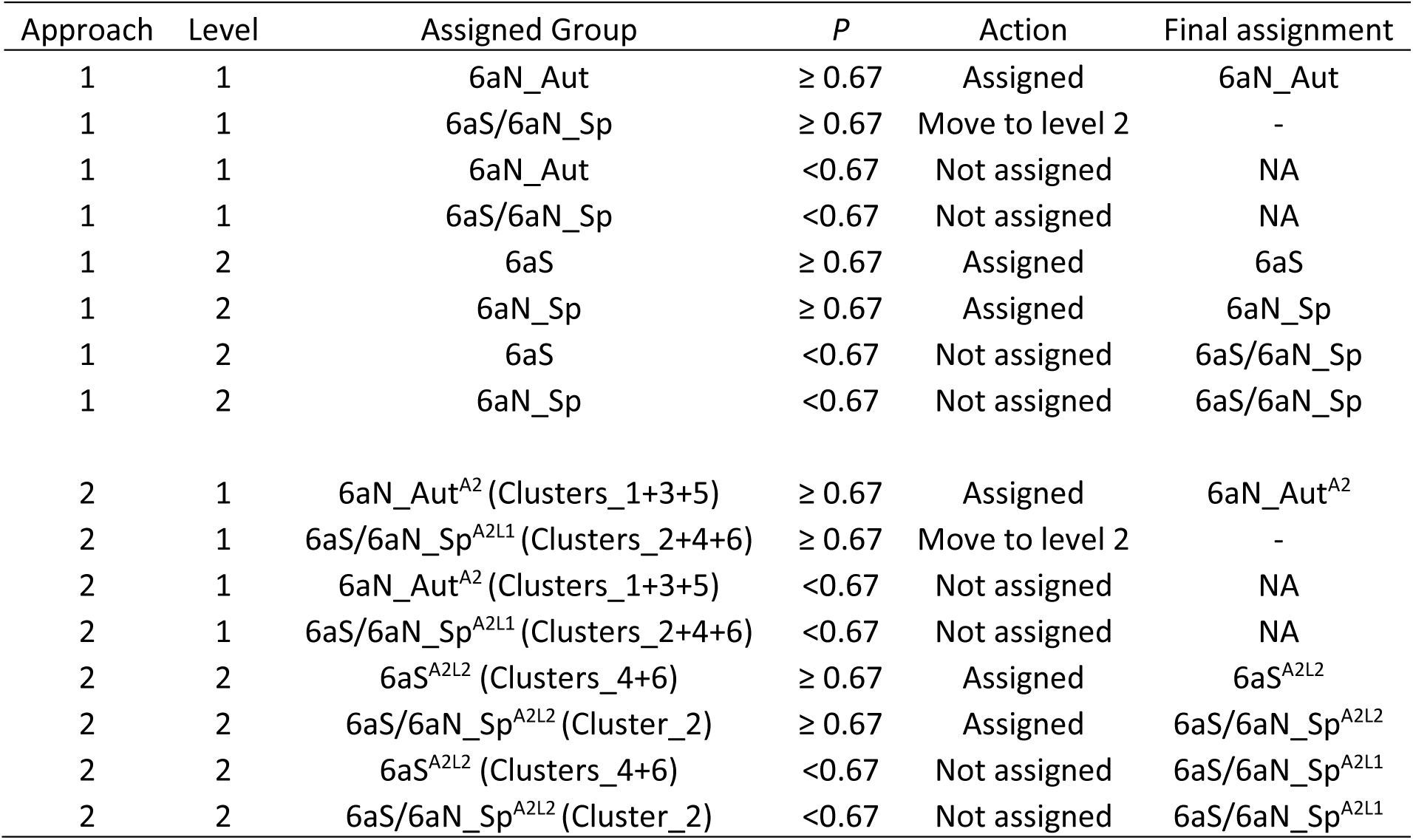
Assignment decision table, indicating the assignment steps in relation to the assignment threshold probability (*P*).

| Approach | Level | Assigned Group | $P$ | Action | Final assignment |
| --- | --- | --- | --- | --- | --- |
| 1 | 1 | 6aN_Aut | $\geq 0.67$ | Assigned | 6aN_Aut |
| 1 | 1 | 6aS/6aN_Sp | $\geq 0.67$ | Move to level 2 | - |
| 1 | 1 | 6aN_Aut | $< 0.67$ | Not assigned | NA |
| 1 | 1 | 6aS/6aN_Sp | $< 0.67$ | Not assigned | NA |
| 1 | 2 | 6aS | $\geq 0.67$ | Assigned | 6aS |
| 1 | 2 | 6aN_Sp | $\geq 0.67$ | Assigned | 6aN_Sp |
| 1 | 2 | 6aS | $< 0.67$ | Not assigned | 6aS/6aN_Sp |
| 1 | 2 | 6aN_Sp | $< 0.67$ | Not assigned | 6aS/6aN_Sp |
| 2 | 1 | 6aN_Aut<sup>A2</sup> (Clusters_1+3+5) | $\geq 0.67$ | Assigned | 6aN_Aut<sup>A2</sup> |
| 2 | 1 | 6aS/6aN_Sp<sup>A2L1</sup> (Clusters_2+4+6) | $\geq 0.67$ | Move to level 2 | - |
| 2 | 1 | 6aN_Aut<sup>A2</sup> (Clusters_1+3+5) | $< 0.67$ | Not assigned | NA |
| 2 | 1 | 6aS/6aN_Sp<sup>A2L1</sup> (Clusters_2+4+6) | $< 0.67$ | Not assigned | NA |
| 2 | 2 | 6aS<sup>A2L2</sup> (Clusters_4+6) | $\geq 0.67$ | Assigned | 6aS<sup>A2L2</sup> |
| 2 | 2 | 6aS/6aN_Sp<sup>A2L2</sup> (Cluster_2) | $\geq 0.67$ | Assigned | 6aS/6aN_Sp<sup>A2L2</sup> |
| 2 | 2 | 6aS<sup>A2L2</sup> (Clusters_4+6) | $< 0.67$ | Not assigned | 6aS/6aN_Sp<sup>A2L1</sup> |
| 2 | 2 | 6aS/6aN_Sp<sup>A2L2</sup> (Cluster_2) | $< 0.67$ | Not assigned | 6aS/6aN_Sp<sup>A2L1</sup> |

The DAPC results supported the previous indications of temporal stability within each of the putative population areas with samples from the same putative populations clustering together (ESM Figure 1). Therefore, the temporal samples were combined to form seven groups (*6aS*, *CS*, *IS*, *6aN_Aut*, *6aN_Sp*, *NS*, *DWN*), which represented the putative populations in the study area. The DAPC and MSA analyses were run again on the pooled samples. Pairwise multi-locus *F_ST_* analyses of the pooled *baseline dataset* indicated significant differentiation between all baseline population groups (Table 1). The lowest level of differentiation was between the *6aN_Aut* and *NS* groups. The level of differentiation between these groups (*F_ST_* = 0.016) was lower than the average differentiation (0.032) between all of the samples within the *6aN_Aut* pool (ESM Table 4). The highest level of differentiation was between the *6aN_Sp* group and the other groups. There was also a very low level of differentiation between the *DWN* group and the *CS* and *IS* groups, whilst the *DWN* group had a high level of differentiation from the *NS* group.

The DAPC results indicated the same pattern of structure as the *F_ST_* analyses (Figure 3) and also as those observed in Han *et al*. (2020) based on whole genome analyses. The highest level of discrimination observed in the DAPC analyses was along the primary axis (74%) and concerned the *6aS* and the *6aN_Aut* groups, though some outliers were evident. The *6aS* and *6aN_Sp* groups were discriminated primarily on the secondary axis (18%). These groups partially overlapped, indicating a lower potential to accurately discriminate between them. The *6aS*, *CS* and *IS* groups overlapped, indicating that the current marker panel cannot be used to distinguish these groups with a high level of accuracy. Therefore, the *CS* and *IS* groups were removed from the baseline data and excluded from further analyses. There is no evidence of significant numbers of herring from these groups being present in Division 6.a (Farrell *et al*., 2021). DAPC also indicated an overlap and an inability to distinguish between the *6aN_Aut* and *NS* groups. There is currently no evidence to support the assertion that the North Sea autumn spawning herring comprise a different population to the *6aN_Aut* herring (Farrell *et al*., 2021), however this distinction was not the focus of the current study, as such the *NS* samples were removed from further analyses. The *DWN* group was confirmed to be distinct from the *NS* group though it could not be reliably discriminated from the *CS* and *IS* groups with the current panel of markers and as such the *DWN* group was removed from further analyses. The resulting reduced *6a baseline dataset* consisted only of the 43 samples from populations that are confirmed as being present in Division 6.a i.e. *6aS*, *6aN_Aut* and *6aN_Sp*.

**Figure 3.**
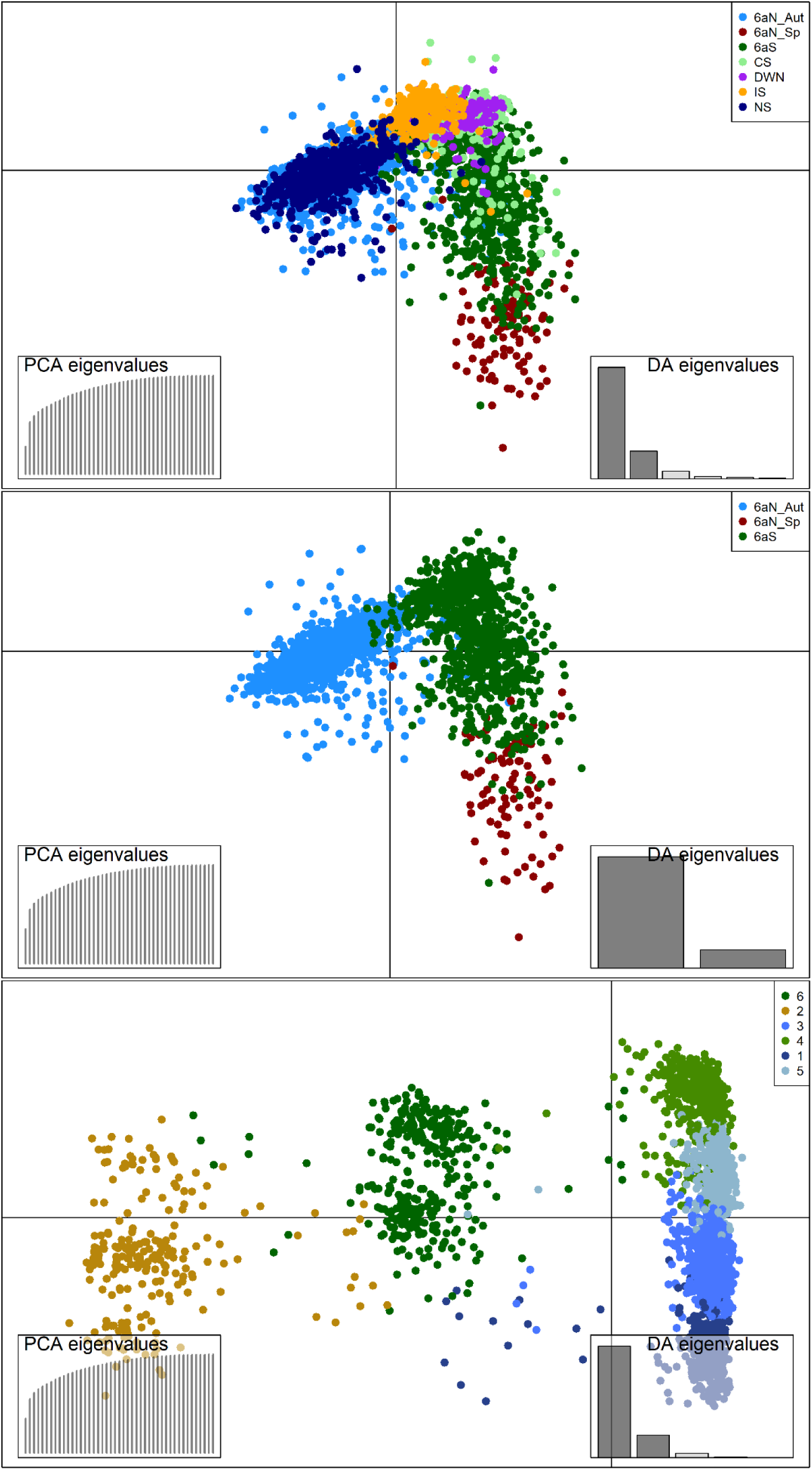
Discriminant Analysis of Principal Components (DAPC) of (top) the pooled *baseline dataset* (middle) the *6a baseline dataset* (bottom) the *clustered 6a baseline dataset*.

Clustering analyses of the *6a baseline dataset* indicated that six clusters were the optimum number to provide the most accurate division of the samples based on their assumed population of origin (ESM Table 5). DAPC of the *clustered 6a baseline dataset* indicated clear division between the clusters with minimal overlap (Figure 3), suggesting that an SVM model-based assignment using this approach would have a high accuracy. The majority of the *6aN_Aut* individuals were represented by the combined *Clusters_1+3+5,* and the majority of the *6aS* individuals by the combined *Clusters_4+6* (Table 2). The majority of *6aN_Sp* individuals were in *Cluster_2*, however this cluster also contained a significant proportion of *6aS* individuals. These individuals were primarily from the samples of late spawning herring collected in Division 6.a.S in January and February. In terms of cluster composition, *Clusters_1+3+5* comprised 98% *6aN_Aut* samples and as such were considered, for the purposes of the assignment model, a proxy for that population group. *Clusters_4+6* comprised 89% *6aS* and 10% *6aN_Aut*. There is some evidence that the *6aN_Aut* individuals in these clusters may be misidentified *6aS* herring or strayers from *6aS* (see Farrell *et al*., 2021), therefore, *Clusters_4+6* were considered to represent *6aS* for the purposes of assignment. *Cluster_2* comprised 54% *6aS* and 44% *6aN_Sp* and was considered, for the purposes of the assignment, to represent a mix of *6aS* and *6aN_Sp* herring. The resulting *clustered dataset* comprised *Clusters_1+3+5*, *Clusters_4+6*, *Cluster_2*. In order to simplify the nomenclature and align it with the *Approach 1* assignment, from this point on these clusters will be referred to as *6aN_Aut^A2^*, *6aS^A2L2^* and *6aS/6aN_Sp^A2L2^*, respectively.

**Table 5.**
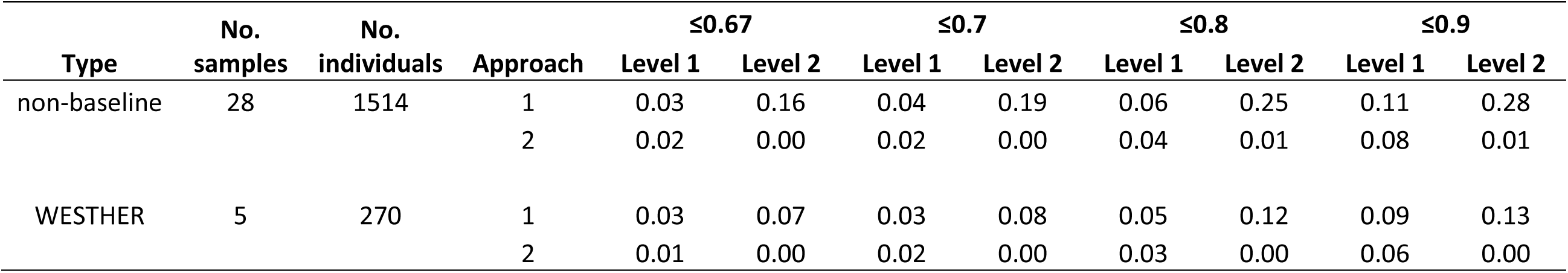
The average proportion of non-baseline and WESTHER samples falling below a range of assignment thresholds for the *Approach 1-Level 1* and *2* and *Approach 2-Level 1* and *2* assignments. The individual sample proportions are in ESM Table 8.

| Type | No.<br>samples | No.<br>individuals | Approach | ≤0.67 |  | ≤0.7 |  | ≤0.8 |  | ≤0.9 |  |
| --- | --- | --- | --- | --- | --- | --- | --- | --- | --- | --- | --- |
|  |  |  |  | Level 1 | Level 2 | Level 1 | Level 2 | Level 1 | Level 2 | Level 1 | Level 2 |
| non-baseline | 28 | 1514 | 1 | 0.03 | 0.16 | 0.04 | 0.19 | 0.06 | 0.25 | 0.11 | 0.28 |
|  |  |  | 2 | 0.02 | 0.00 | 0.02 | 0.00 | 0.04 | 0.01 | 0.08 | 0.01 |
| WESTHER | 5 | 270 | 1 | 0.03 | 0.07 | 0.03 | 0.08 | 0.05 | 0.12 | 0.09 | 0.13 |
|  |  |  | 2 | 0.01 | 0.00 | 0.02 | 0.00 | 0.03 | 0.00 | 0.06 | 0.00 |

### Assignment model development

The optimum numbers of PCs for the *Approach 1-Level 1* dataset and *Approach 1-Level 2* dataset, determined as the values with the lowest root mean squared error (RMSE) following DAPC cross-validation, were 40 and 35, respectively. The optimum number of PCs for the *Approach 2-Level 1* dataset and *Approach 2-Level 2* dataset were 30 and 5, respectively. There was however little difference between the number of PCs retained in all cases suggesting that the assignment is not sensitive to this parameter. The optimum model and kernel for the assignment model were the SVM model and the radial basis function (RBF) kernel. Grid search indicated the optimum values for cost and gamma in *Approach 1-Level 1* and *Approach 2-Level 1* were 1 and 0.33, respectively and in *Approach 1-Level 2* and *Approach 2-Level 2* were 1 and 0.5, respectively.

There was little difference between the self-assignment accuracy of *Approach 1-Level 1* and *Approach 2-Level 1* (ESM Figures 2 & 3 and Table 3). Both approaches resulted in self-assignment rates greater than 90% and neither approach was observed to be particularly sensitive to the number of individuals in the training data. Similarly, neither approach was observed to be particularly sensitive to the proportion of highest *F_ST_* loci used in the analyses. The main difference between the two approaches at *Level 1* was the higher probabilities of assignment and lower error observed in the *K*-fold analyses in *Approach 2* (ESM Figure 3 and Table 3). Conversely there were large differences between the two approaches in the *Level 2* assignments, where *Approach 1-Level 2* did not confidently assign *6aN_Sp* samples to their baseline, whereas *Approach 2-Level 2* achieved near perfect self-assignment to both the *6aS^A2L2^* and *6aS/6aN_Sp^A2L2^*.

*The Approach 1-Level 1* assignment was more sensitive to the number of loci *than Approach 1-Level 2* (ESM Figure 4 and ESM Table 6). This was particularly notable for the *6aS/6aN_Sp* group in *Level 1*, where there was a significant drop in assignment accuracy and an increase in the number of outliers below 50% of loci. This indicated that at least twenty-three of the forty-five loci were required for accurate assignment at this level. Ideally over 60% (27 loci) should be genotyped at this level to ensure assignment accuracy. The *Approach 1-Level 2* assignment was not very sensitive to the number of loci and there was little difference in the accuracy of assignment down to 20% of the loci. *The Approach 2-Level 1* assignments had a similar pattern of sensitivity to the number of loci as the *Approach 1* assignments indicating that a minimum of 60% of loci should be genotyped at this level to ensure assignment accuracy. The *Approach 2-Level 2* assignments had a higher level of accuracy than the *Approach 1-Level 2* assignments and again were less sensitive to the number of markers than the *Level 1* assignments.

**Figure 4.**
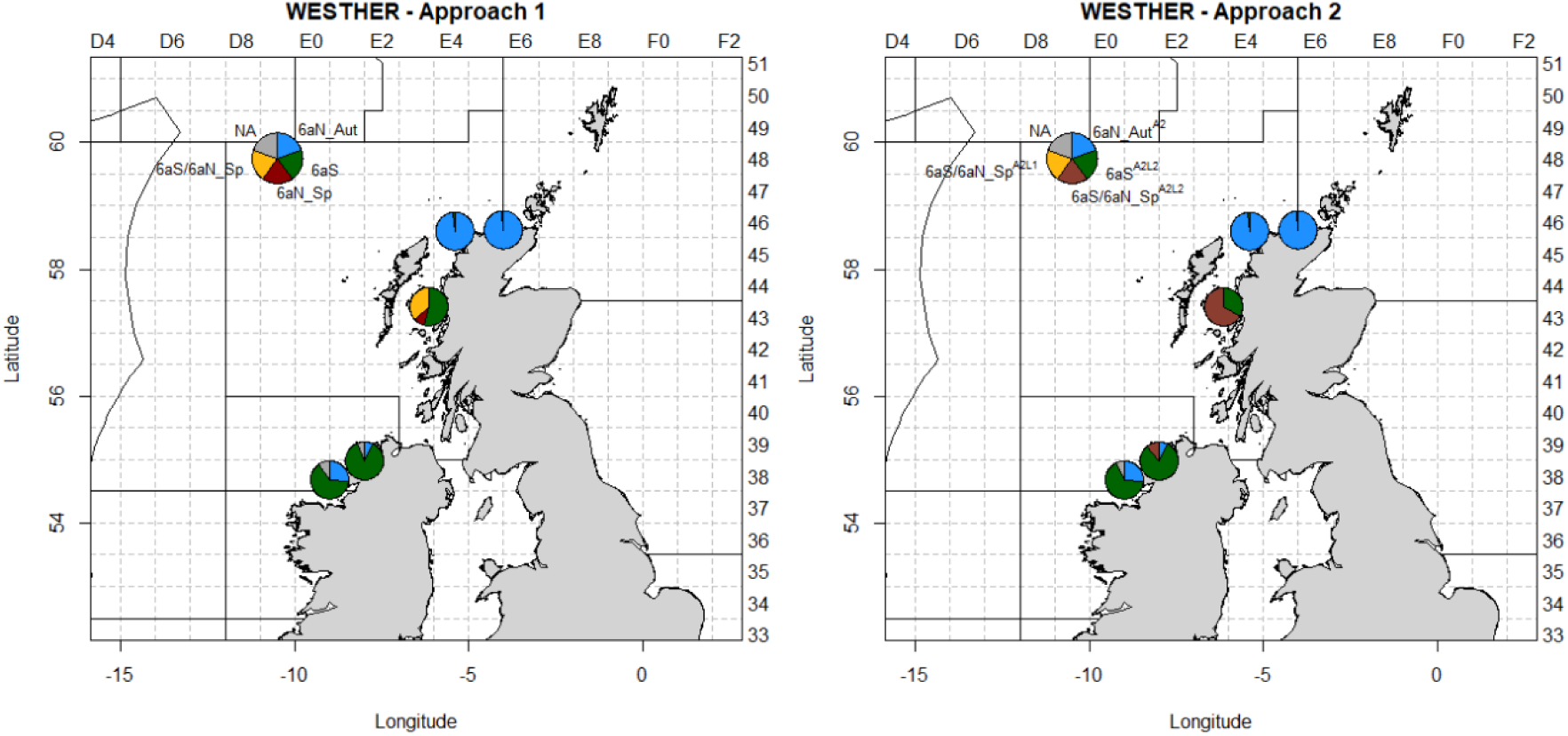
Assignment output of the archive samples from the WESTHER project following (left) Approach 1 and (right) Approach 2.

### Assignment model validation with archive samples

The final assignment call of each individual in the archive WESTHER samples was based on a combination of the *Level 1* and *Level 2* assignments according to the assignment decision table (Table 4). The assignments of the WESTHER *6aN_Aut* samples from 2003 (*6aN_03*) and 2004 (*6aN_04*) displayed near perfect assignment to the *6aN_Aut* groups, with only one individual in each of the years being misassigned to the *6aS* groups (Figure 4 and ESM Table 7). The assignment of the 2003 *6aS* WESTHER sample (*6aS_03*) was more uncertain, with a quarter of individuals misassigned to the *6aN_Aut* groups (ESM Table 7). The 2004 *6aS* sample (*6aS_04*) had a higher level of correct assignment (>80%). The assignment of the 2004 *6aN_Sp* (*6aN_Sp_04*) sample indicated perfect assignment, at *Level 1* in both approaches, to the *6aS/6aN_Sp* groups. As expected, the *Approach 1-Level 2* assignment displayed a high rate of misassignment (53%) to the *6aS* group and a high rate of below threshold individuals (37%), which could not be confidently split below the level of *6aS/6aN_Sp*. The *Approach 2-Level 2* assignment provided a more robust assignment with 67% of individuals assigned to *6aS/6aN_Sp^A2L2^*. It should be noted that the *6aN_Sp_04* WESTHER sample did not fulfil the criteria of baseline samples as defined in the current study as 27% of the individuals were maturity stage 2 individuals (ESM Table 2).

The average proportion of unassigned individuals in the WESTHER samples increased with increasing assignment threshold for *Approach 1-Level 1* and *2* but only for *Level 1* in *Approach 2* (Table 5). Analysis of the individual rather than average values (ESM Table 8) showed differences between the individual samples. All of the *6aN_Aut* samples’ individual assignments had a probability greater than 0.9, indicating they were at least nine times more likely that the alternate assignment (ESM Table 8). The assignment probabilities for the 2003 *6aS* individuals were more variable at *Level 1* for *Approach 1* and *2*, highlighting a degree of uncertainty around the assignment of some of the individuals, however the *Level 2* assignments all had a probability greater than 0.9 (ESM Table 8). The 2004 *6aS* individuals also showed some uncertainty at *Level 1* and at *Approach 1-Level 2*. *The Approach 2-Level 2* assignments all had a probability greater than 0.9. The individuals of the *6aN_Sp_04* sample had the highest proportion of unassigned individuals at the higher thresholds for *Approach 1-Level 2*. *The Approach 2-Level 2* assignments all had a probability greater than 0.9.

### Exploratory analyses with contemporary non-baseline samples

The quarter 1 non-baseline samples from Division 6.a.N came primarily from the Scottish West Coast International Bottom Trawl Survey (SWC-IBTS) and comprised a number of small samples of herring of mixed maturity stages (Figure 5 and ESM Tables 2 & 3). The most northerly samples (*6aN_Sp_19a* and *6aN_Sp_19b*) were dominated by resting (*Stage 5*) individuals and the assignments indicated a significant proportion of *6aN_Aut* individuals, though assignments to both the *6aS* and *6aS/6aN_Sp* groups were also evident (ESM Table 7). One haul to the north of the Hebrides (*6aN_Sp_19d*) comprised primarily maturing (*Stage 2*) and spawning (*Stage 3*) herring that assigned to the *6aS* and *6aS/6aN_Sp* groups. In the south Minch area there was a significant proportion of immature juvenile (*Stage 1*) individuals in one sample (*6aN_Sp_19h*), which assigned primarily to *6aS*. The other two hauls in this area (*6aN_Sp_19f* and *6aN_Sp_19g*) also had the majority of assignments to the *6aS/6aN_Sp* groups.

**Figure 5.**
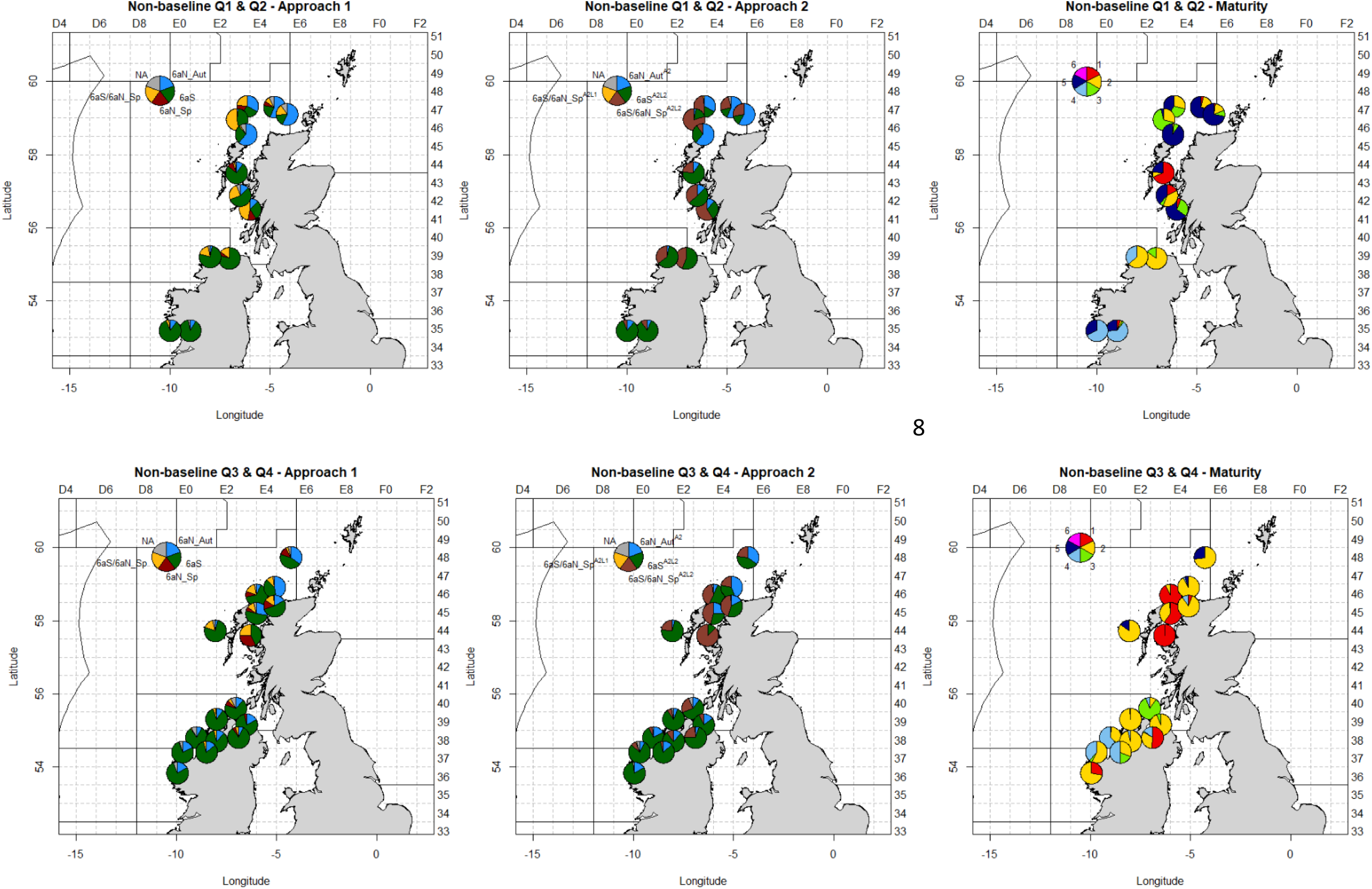
The assignment outputs and maturity stages of the contemporary non-baseline samples divided by quarter. Note the exact catch positions have been adjusted to minimise the overlap of the pie charts.

The quarter 1 and 2 samples from Irish coastal waters were collected from bycatch of commercial vessels and comprised four samples, one sample from Lough Foyle (*6aS_19b*), one from Lough Swilly (*6aS_19a*) and two from Galway Bay (*6aS_18b* and *6aS_18c*) (Figure 5 and ESM Table 2). The Lough Foyle sample contained mainly *Stage 2* individuals with a small proportion of *Stage 3* individuals and the Lough Swilly sample, which was caught two days earlier, contained predominately *Stage 2* individuals and some spent (*Stage 4*) fish. The assignment of both samples was quite similar with the majority of individuals assigned to *6aS* and most of the remaining samples to *6aS/6aN_Sp*. The Galway Bay samples were caught later in quarter 1 and 2 and had similar maturity staging with primarily *Stage 4* and *Stage 5* individuals. The assignments of the two samples were also quite consistent with each other with the majority of individuals assigned to *6aS*.

The quarter 3 non-baseline samples all came from Division 6.a.N and comprised samples from acoustic survey and monitoring fishery catches (see Mackinson *et al*, 2019a; 2019b; 2021). The maturity stages and length frequencies of the samples were notably different to the quarter 1 samples as there was a significant proportion of *Stage 1* and *2* fish (Figure 5 and ESM Tables 2 & 3). The three samples from the Minch (*6aN_18a, 6aN_19h* and *6aN_19i*) primarily comprised *Stage 1* individuals, which assigned mainly to *6aS* and *6aS/6aN_Sp*. The two samples from northwest of Cape Wrath (*6aN_18b* and *6aN_19e*) were composed of mainly *Stage 2* individuals and comprised a mix of *6aN_Aut*, *6aS* and *6aS/6aN_Sp*. The two samples from July (*6aN_19f* and *6aN_19g*), caught west of the Hebrides and north of Scotland, had similar maturities with mainly *Stage 2* individuals and a smaller proportion of *Stage 5* fish. The majority of individuals in the west of Hebrides sample were assigned to *6aS*. Conversely the north of Scotland sample, which was caught adjacent to the 4°W line of longitude, contained a significant proportion of *6aN_Aut* fish in addition to significant proportions of *6aS* and 6aS/*6aN_Sp* individuals.

The quarter 4 non-baseline samples all came from Divisions 6.a.S, 7.b and Lough Foyle and comprised samples from monitoring fishery catches and bycatch. The nine samples, caught over 5 years, contained a wide range of maturity stages, from *Stage 1* to *Stage 4* yet the assignments were relatively consistent across all of the samples (Figure 5 and ESM Tables 2 & 7). The five Division 6.a.S samples contained mainly a mixture of *Stage 2* and *Stage 4* individuals with a small number of *Stage 3* fish and the majority of individuals were assigned to *6aS* in all samples (ESM Table 7). The Division 7.b sample (*6aS_19d*) contained *Stage 1* and *Stage 2* fish and the assignments followed the same pattern as the Division 6.a.S samples. The three Lough Foyle samples were collected in three different years and had markedly different maturity stages, with one sample (*6aS_17f*) dominated by Stage 3 fish, one sample (*6aS_18e*) predominately *Stage 2* and the third sample (*6aS_20b*) a mixture of *Stages 1-4* fish (Figure 5 and ESM Table 2). Regardless, the assignment outputs were broadly similar between the three samples and with the Division 6.a.S, 7.b samples, apart from a higher proportion of *6aS/6aN_Sp* fish in two of the samples (ESM Table 7).

Similar to the WESTHER samples, the average proportion of unassigned individuals increased significantly with increasing assignment threshold for *Approach 1-Level 1* and in particular for *Approach 1-Level 2* (Table 5). This was not as significant in *Approach 2*, where there was a minor increase in the proportion of unassigned individuals at increasing *Level 1* thresholds and almost no increase in the *Level 2* assignments. Analysis of the individual rather than average values (ESM Table 8) showed differences between the individual samples. The non-baseline quarter 1 samples from Division 6.a.N displayed the highest level of unassigned individuals, particularly at the *Approach 1- Level 2* assignment. The *Approach 2* assignment of the majority of samples resulted in a lower incidence of unassigned individuals at all threshold levels.

## Discussion

The genetic markers and assignment methods presented in the current study constitute a ‘tool box’ that can be used for the assignment of herring caught in mixed survey and commercial catches in Division 6.a into their population of origin with a high level of accuracy (>90%). This will enable the population assignment of commercial catch and acoustic survey (e.g. MSHAS) samples, which will facilitate the development of separate stock assessments for the populations in this area.

Both assignment approaches had a high level of self-assignment accuracy though it was notable that there was a higher power to discriminate between the groups in *Level 1* than the groups in *Level 2*. In *Approach 1-Level 2* there was a weakness in the model in discriminating between the *6aN_Sp* samples and the quarter 1 late-spawning *6aS* samples. This was due in part to the small number of samples of *6aN_Sp* in particular, and potentially also to the genetic markers used not being optimised for distinguishing between these groups. There are inherent difficulties in sampling these groups for which no specific fishery currently exists and which spawn in areas that are subject to unfavourable weather conditions for sampling at the time of spawning. Further sampling of both groups is required though, and efforts are also underway to conduct WGS on representative samples from these groups to identify more informative markers. In any case *Approach 2* mitigated this issue by combining them into a single group (*6aS/6aN_Sp^A2L2^ (Cluster_2)*) for assignment purposes, which resulted in higher classification success and lower rate of unassigned individuals. In this case the majority of *6aS* fish could be separated from the other two populations with a high level of accuracy and only the minority of *6aS* fish were left in an unsorted mix with *6aN_Sp*. This is considered preferable to the outcome of *Approach 1*, as for the purposes of splitting catch and survey samples it is better to be able to assign some level of grouping rather than have a high abundance of unassigned individuals. Mixed categories such as *6aS/6aN_Sp^A2L2^* can be acknowledged in the overall abundance estimates but retained in a separate category that is not allocated to a single stock, which can also act as a precautionary buffer for any undetected misassignments. In each assignment approach at least 60% of the 45 markers were required to ensure accurate self-assignment, which indicates that there is a level of redundancy built into the panel of markers as was expected given that the markers are distributed among fourteen loci. This redundancy is an advantage when analysing unknown samples as it allows up to 40% missing data in the genotypes of individuals. Missing genotypes may occur when analysing suboptimal quality samples collected from commercial catches or when analysing older archive samples, which have not been stored under optimum conditions.

The genetic assignment of the archive WESTHER samples confirmed longer term temporal stability of the SNP panel in the Division 6.a populations over a period of at least eighteen spawning seasons, which is a temporally relevant time scale for the purposes of stock assessment. Whilst the 2003 and 2004 *6aN_Aut* WESTHER samples assigned near perfectly to the *6aN_Aut* groups, the assignments of the 2003 and 2004 *6aS* WESTHER samples were not as confident. There were a significant number of mis-assignments to *6aN_Aut* groups, particularly in the 2003 *6aS* sample. This spawning sample was collected in October which is earlier than any of the contemporary samples in the *6a baseline dataset* and it is possible that these misassigned individuals shared some genetic similarities related to spawning time with the *6aN_Aut* autumn spawning herring. Historically autumn spawning herring were abundant in Division 6.a.S and particularly in Division 7.b where they supported local fisheries (see Farrell *et al*., 2021; ICES, 2015) however no autumn spawning was observed or sampled in this area in the course of the current study (i.e. since 2014). In fact, no spawning baseline samples were collected in Division 7.b throughout the study despite repeated sampling attempts, suggesting that either spawning in that area is at a very low level or has not occurred in recent years. However, the non-spawning herring caught in Division 7.b, genetically assigned with a high probability to *6aS*. Continued efforts should be made to ensure any spawning activity in Division 7.b is sampled if it occurs and the data added to the baselines.

The assignment of the non-baseline samples also provided an additional layer of validation of the assignment model and an interesting exploratory analysis of potential mixing of the different populations in Division 6.a. The Division 6.a.S and 7.b sample assignments were relatively consistent across all quarters indicating stability in the composition of herring shoals in the area. In all samples a minority proportion of individuals were assigned to *6aN_Aut* though this was mostly in keeping with the expected error rate of the assignment model, which was higher for *6aS* and *6aS/6aN_Sp* than for *6aN_Aut*. The samples from Lough Foyle were shown to be genetically and biologically the same as the *6aS* samples underlining the inappropriateness of the existing classification of Lough Foyle as part of the *6aN_Aut* stock.

Whilst there was consistency in the assignment of the samples collected in Division 6.a.S and 7.b, the assignment of those from Division 6.a.N (excluding Lough Foyle) indicated a significant degree of mixing of different populations. *6aS* and mixed *6aS/6aN_Sp* herring comprised a varying but significant proportion of all samples and were far in excess of the expected error rate. The assignment of the juvenile samples from the Minch primarily to *6aS*, *6aN_Sp*, *6aS/6aN_Sp* instead of *6aN_Aut* was not unexpected given the existing knowledge about the larval drift in the area and the lack of differentiation between the *6aN_Aut* and North Sea autumn spawning herring. It is well documented that the larvae of autumn spawning herring off the northwest of Scotland are carried in easterly flowing currents and spend their juvenile phase in the North Sea (Heath, 1989; Heath, 1990; Heath *et al*., 1987; MacKenzie, 1985; Saville and Morrison, 1973). The mixed nature of the samples collected off Cape Wrath during the spawning season for *6aN_Aut* herring indicated a need for ongoing monitoring of survey and commercial catches in this area as any future fisheries in this area will likely be mixed stock fisheries. The presence of *6aN_Sp* herring in the samples was also of interest as this population used to be the dominant population in the region but was reported to have collapsed in the 1950s (Baxter, 1958). Despite this, spring spawning herring were still known to comprise up to 38% of the catches off the north of Scotland, west of the 4°W line of longitude and in the North Minch in the 1960’s (Saville, 1970). However, as the autumn spawning component was more abundant, the newly developing stock assessments at the time were restricted to that group and the spring spawning herring were not distinguished, which over time led to them being merged with the autumn spawners for assessment purposes. The results of the current study suggest the spring spawners are still present in the area. It is not currently possible to separate them from the late spawning *6aS* herring so no conclusions can be drawn about their relative abundance, but further efforts should be directed towards improving the sampling of this population. The two samples collected in July west of the Hebrides and North of Scotland also offer some insight into the future assignments of the MSHAS samples that are collected during this period. The west of Hebrides sample comprised primarily *6aS* individuals with a smaller proportion of *6aS/6aN_Sp* fish (Figure 5), whilst the north of Scotland sample, which was caught adjacent to the 4°W line of longitude, contained a significant proportion of *6aN_Aut* fish in addition to significant proportions of *6aS*, *6aN_Sp* and *6aS/6aN_Sp* individuals. Therefore, the current approach of splitting the MSHAS data using the 56°N line of latitude and the 7°W line of longitude to delineate the Division 6.a stocks is inappropriate and should be replaced with the genetic assignment approach.

One weakness of the assignment model in the current study is that it is solely based on the populations empirically proven to occur within Division 6.a (i.e. *6aS*, *6aN_Aut* and *6aN_Sp*) and does not include adjacent populations. The initial genetic analyses of the *full baseline dataset* in the current study and those in Han *et al*. (2020) demonstrate that the Irish Sea herring and the Celtic Sea herring are distinct from each other and from the other populations in Division 6.a. However, they are genetically closely related to the herring in Division 6.a.S and as such it is difficult to distinguish them with a high degree of certainty using the current marker panel. Inclusion of these populations in the baseline dataset would increase the overall uncertainty of the assignments. Despite the assertions of the WESTHER project (Hatfield *et al*., 2005) there is no definitive evidence that a significant abundance of herring from either of these populations migrate to Division 6.a (see Farrell *et al*., 2021). Therefore, their inclusion in the baseline datasets is not warranted at this time. The WESTHER project provided an illustration of the dangers of including multiple populations in a baseline when the power of discrimination between the populations is low. The inevitable outcome is that mixed samples will be weakly assigned and will have a high rate of misassignment. This can lead to the incorrect conclusion that mixed samples come from a larger number of source populations when the converse may be true. In the current study there is the potential to misassign individuals from the Celtic Sea and Irish Sea populations, if they were present in Division 6.a, however the assignment in its current form is still a significant improvement on the existing method of splitting the stocks based on geographic delineation, which is proven to be inappropriate. Efforts should be made to identify further population specific genetic markers that may increase the discriminatory power between closely related populations. For this reason, the current marker panel should be considered the best available at the current time, but continued efforts should be made to develop it further.

The current study has also highlighted some of the potential stock identification issues that are apparent with the North Sea herring. The lack of differentiation between the *6aN_Aut* herring and the North Sea autumn spawning herring suggests that the 4°W line of longitude is also inappropriate as a stock delineator. Though this has been recognised since its inception, as Saville and Bailey (1980) noted, ‘*the dividing line between VIa and the North Sea (sub-area IV) at 4°W longitude was not chosen on any criterion of herring stock differentiation but for convenience in statistics collection*’. Further, the current study has demonstrated that *6aS* herring may be found up to at least the 4°W line of longitude and Farrell *et al*. (2021) demonstrated the uncertainty in the composition of HERAS hauls in close proximity east of this line. The winter spawning Downs herring have also been shown to be a distinct and separate population to the North Sea autumn spawning herring and are relatively easily distinguished with the genetic markers in the current panel. The extent of the distribution of Downs herring in the North Sea area and their abundance in the HERAS or in the commercial catches in Divisions 4.a and 4.b are currently unknown. There are also known and demonstrated issues of mixing of the North Sea autumn spawning herring with Western Baltic herring to the east (Bekkevold et al., In preparation) and with Norwegian Spring Spawning herring to the North (Berg et al., 2017). What is required now is a cohesive effort to study all of these stock identification issues, and those in the current study, together rather than treating them all separately. The ideal scenario may be to develop a universal marker panel that can discriminate all of the populations that could potentially be surveyed or caught in the Northeast Atlantic area (FAO Major Fishing Area 27). In theory this would solve some of the issues outlined above, it would, however, also create a cost-benefit issue relating to the use of more extensive and expensive panels. In order to differentiate a wider range of populations, including those in the Baltic Sea, the panel would certainly need to comprise a larger number of genetic markers. The markers that may be suitable for discriminating between some of the Baltic Sea populations would likely not be informative for the populations west of Ireland and Britain (see Han *et al*., 2020). Therefore, using a universal panel of markers on a sample caught to the west of Ireland and Britain, which is highly unlikely to contain any Baltic herring, would represent a degree of wasted resources. If the universal panel was used on a sample caught in the eastern North Sea, then the presence of the Baltic markers may actually be beneficial as there is the potential for some Baltic Sea populations to be present in this area. The difficulty arises in defining the cut-off points on where the use of the universal panel is justified and where it is wasteful. Such a definition is akin to delineating stocks based on geographic or statistical areas such as ICES Divisions and inevitably introduces an element of subjectivity that may bias the results. It also introduces issues concerning the temporal stability of such definitions in an era of changing environmental conditions and documented changes in species distributions. Therefore, the use of a universal or specific marker panel is a topic that requires very careful consideration and rigorous empirical testing.

Implementation of a universal marker panel would also necessitate the development and implementation of a standard assignment approach across multiple jurisdictions. The use of the SVM model in *assignPOP* in the current study was favoured over traditional methods that rely on genotypic frequency distribution, as it was not constrained by underlying assumptions of HWE and linkage equilibrium. Further it enabled a transparent and reproducible approach that could be clearly understood by non-geneticists, which is important if genetic stock identification methods are to be accepted and widely implemented as part of standard fisheries data collection protocols. Though not used in the current study, *assignPOP* also allows for the use of non-genetic markers either as standalone assignment models or in combined models with genetic data. This option may be attractive to institutes that have long time series of morphometric and meristic based stock identification data as the transition to genetic based methods can be made easier with direct comparison of the different data sets that are capable of being conducted within the same analyses.

The SNP panel deployed in the current study was composed of adaptive markers that are known to be under diversifying selection and proven to be associated with local ecological adaptation (Han *et al*., 2020; Martinez Barrio *et al*., 2016). Genetic markers associated with loci under selection have been proven to provide better resolution to distinguish population structure in herring than neutral genetic markers (Bekkevold *et al*., 2016; Han *et al*., 2020). However, such high-graded adaptive markers may undergo more rapid changes in allele frequencies within populations than putatively neutral genetic markers, particularly in situations of dynamic environmental conditions (Jorde *et al*., 2018; Nielsen *et al*., 2012). In the current study the contemporary baseline spawning samples collected from 2014 to 2021 (seven spawning seasons) indicated temporal stability of the genetic markers within the different populations. Thus, these SNPs were appropriate for the purposes of stock identification in the current study. However, it is advisable to continue to collect and analyse baseline spawning samples regularly to monitor any changes in allele frequencies within the populations in the assignment model in order to prevent erroneous assignments of mixed samples. This also raises the question of how long to retain samples in the baseline dataset. As populations evolve and respond to changing conditions, older samples may become less relevant as baseline samples and may not represent the populations in their current state, which may have a negative impact on the assignments. Thus baseline samples should perhaps be limited to a time scale that is relevant to the current population e.g. the longevity of the species. Given the relatively short history of effective genetic stock identification this has not been an issue to date but should be considered now as genetic stock identification starts to become an important part of marine fish stock assessment.

To date there are few examples of genetic stock assignment being used for the regular assignment of survey or catch data of marine fish into population of origin for the purposes of stock assessment.

These methods have primarily been used for one off studies, that at best have been used to inform management but few have been developed for regular monitoring and data collection (Reiss *et al*., 2009; Waples *et al*., 2008). Genetic stock identification methods have been most commonly used for salmonids, including Atlantic Salmon, *Salmo salar* (Gilbey *et al*., 2016) and species of pacific salmon including coho salmon, *Oncorhynchus kisutch* (Beacham *et al*., 2020). In these studies, self-assignment accuracies of 70-80% were concluded to be acceptable levels of accuracy. The high level of self-assignment accuracy in the current study (>90%) exceeds this and is maintained even in the event of a considerable number of missing genotypes per individual.

If implemented as part of regular data collection on the MSHAS, the genetic stock identification method in the current study will enable the splitting of the survey indices into their constituent Division 6.a populations, which has not previously been possible. As a result, it will be possible to develop a separate stock assessment for the Division 6.a.S, 7.b-c stock. Although, it should be noted that as there were no spawning herring observed or sampled in Divisions 7.b and 7.c, it was not possible to test the assumption that the herring that spawn in these Divisions are the same population as the *6aS* herring. The apparent lack of differentiation between the *6aN_Aut* herring and the North Sea autumn spawning herring also raises the question of whether it is appropriate to conduct a stand- alone assessment on the *6aN_Aut* herring or whether it should be combined with the North Sea autumn spawning herring assessment, though it is beyond the remit of the current study to make this recommendation. What is clear is that the results of the current study have improved the capacity to delineate, survey and assess the herring stocks in Division 6.a and there is a need now to translate this into improved management.

## Supporting information

Farrell_et_al_6a_herring_baseline_ESM_tables_figures

## Data accessibility

All data are provided in the manuscript and extra supplementary tables (ESM Table 1 – ESM Table 8) and figures (ESM Figure 1 – ESM Figure 4) are provided in the file *Farrell_et_al_6a_herring_baseline_ESM_tables_figures.xlsx*.

## Research Ethics, Animal Ethics

No ethical approval was required. All samples were collected opportunistically from fish caught during fisheries surveys and commercial fisheries. No fish were killed for the purpose of the study and no licences were required.

## Author’s contributions

Conceptualisation: EDF, JC, MWC, SML, CN. Data Curation: EDF, SML, SM, CN, SO’C, MO’M, MP, EW. Formal analysis: EDF, LA, DB, NC, JC, MWC, AE, AF, MG, SML, SM, CN, SO’C, MO’M, MP, MEP, EW.

Funding acquisition: EDF, NC, JC, MWC, SML, SM, CN, MP. Investigation: EDF, LA, DB, NC, JC, MWC, AE, AF, MG, SML, SM, CN, SO’C, MO’M, MP, MEP, EW. Methodology: EDF, LA, DB, JC, AE, AF, MG, SML, SM, CN, SO’C, MO’M, MP, MEP, EW. Project administration: EDF, NC, JC, MWC, MG, SML, SM, CN, MP. Resources: EDF, LA, DB, MWC, SML, SM, CN, SO’C, MO’M, MP, MEP, EW. Validation: EDF, LA, DB, NC, JC, MWC, AE, AF, MG, SML, SM, CN, SO’C, MO’M, MP, MEP, EW. Visualisation: EDF, EW.

Writing-original draft: EDF, EW. Writing-review & editing: EDF, LA, DB, NC, JC, MWC, AE, AF, MG, SML, SM, CN, SO’C, MO’M, MP, MEP, EW.

## Acknowledgements

The work in the current study was part funded (2018-2020) by the European Commission’s Executive Agency for Small and Medium-sized Enterprises (EASME) under Service Contract EASME/EMFF/2017/1.3.2.1/SI2.767459. The authors would also like to thank the Killybegs Fishermen’s Organisation, Pelagic Freezer Trawler Association, Scottish Pelagic Fishermen’s Association, Marine Institute, Marine Scotland Science, Pelagic Advisory Council and the Northern Pelagic Working Group of the European Association of Fish Producers Organisations for continued financial and ancillary support throughout the duration of the project (2015-2022). We are also grateful to all the scientists and crew of the RVs ‘*Celtic Explorer*’, ‘*Scotia*’ and ‘*Corystes*’ and all of the commercial vessels and scientists that have helped with the collection of samples. We acknowledge the help and support provided by Kuan-Yu “Alex” Chen with the turning of the assignPOP model. Finally, we acknowledge the valuable contribution of the GENSINC project (GENetic adaptations underlying population Structure IN herring; Research Council of Norway project 254774), which identified the informative markers that made the current study possible.

